# Aurora kinase A mediated phosphorylation of mPOU is critical for skeletal muscle differentiation

**DOI:** 10.1101/589754

**Authors:** Dhanasekaran Karthigeyan, Arnab Bose, Ramachandran Boopathi, Vinay Jaya Rao, Hiroki Shima, Narendra Bharathy, Kazuhiko Igarashi, Reshma Taneja, Tapas K. Kundu

## Abstract

Aurora kinases are Ser/Thr-directed protein kinases which play pivotal roles in mitosis. Recent evidences highlight the importance of these kinases in non-mitotic biological events like skeletal myogenesis. Our earlier study identified POU6F1 (or mPOU) as a novel Aurora kinase A (AurkA) substrate. Here, we report that AurkA phosphorylates POU6F1 at Ser197 and inhibits its DNA binding ability. Delving into POU6F1 physiology, we find that the phospho-mimic (S197D) POU6F1 mutant exhibits enhancement, while wild type (WT) or phospho-deficient (S197A) mutant shows retardation in C2C12 myoblast differentiation. Interestingly, POU6F1 depletion phenocopies S197D-POU6F1 overexpression in the differentiation context. Collectively, our results signify mPOU as a negative regulator of skeletal muscle differentiation and strengthens the importance of AurkA in skeletal myogenesis.

## Introduction

Regeneration of adult skeletal muscle tissue relies on a group of unipotent, mono-nucleated muscle precursor cells called satellite cells (SCs). SCs are quiescent in nature and exhibit a reversible G_0_ phase arrest (1). Injury to myofibrils induce cell cycle re-entry of the resident SCs to give rise to a population of immature myoblasts which eventually proliferate and subsequently undergo terminal differentiation into elongated, multi-nucleated myofibres, an event which observes myoblast fusion. A subset of the myoblasts, however, re-populate the SC niche. Terminal differentiation not only demands hierarchical regulation of muscle specific gene expression mediated by a set of basic helix-loop-helix myogenic regulatory factors (MyoD, Myf5, myogenin and MRF4) but also a permanent exit from the cell cycle (2–4). Accordingly, various regulators like p21, pRb or Aurora kinases have been implicated in the process of differentiation (5–7).

Our previous *in vitro* high-throughput screening had led to the identification of POU6F1, a POU domain containing transcription factor variously known as mPOU, Brn-5, Emb or TCFβ1, as a novel AurkA substrate (7). The POU (<u>*P*</u>it 1, <u>*O*</u>ct 1, and Oct 2, <u>*U*</u>nc 86) domain is a bipartite DNA-binding domain, consisting of two highly conserved regions (POU-specific domain and POU homeodomain) tethered by a variable linker (8–10). The POU domain containing proteins are sub-divided into six broad classes and the members are heavily involved in various developmental processes (11,12). POU6F1 (Fig.1A) is a member of the class VI-POU domain transcription factors and exhibits preferential binding to a variant of the DNA octamer motif (5’-ATGATAAT-3’) (13,14). Although it displays tissue-restricted expression (13), the exact function carried out by this protein is largely unclear.

**Figure 1:**
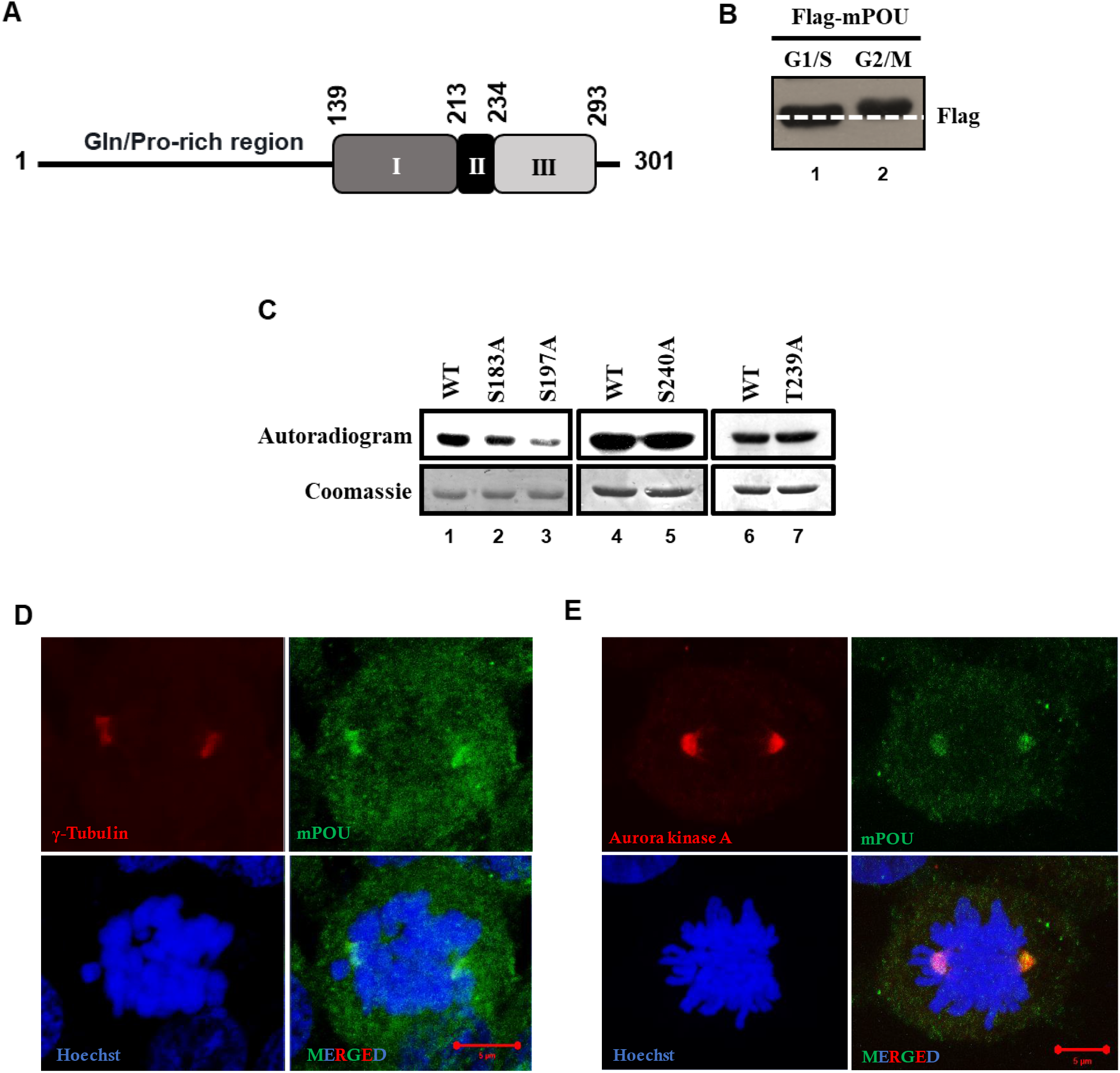
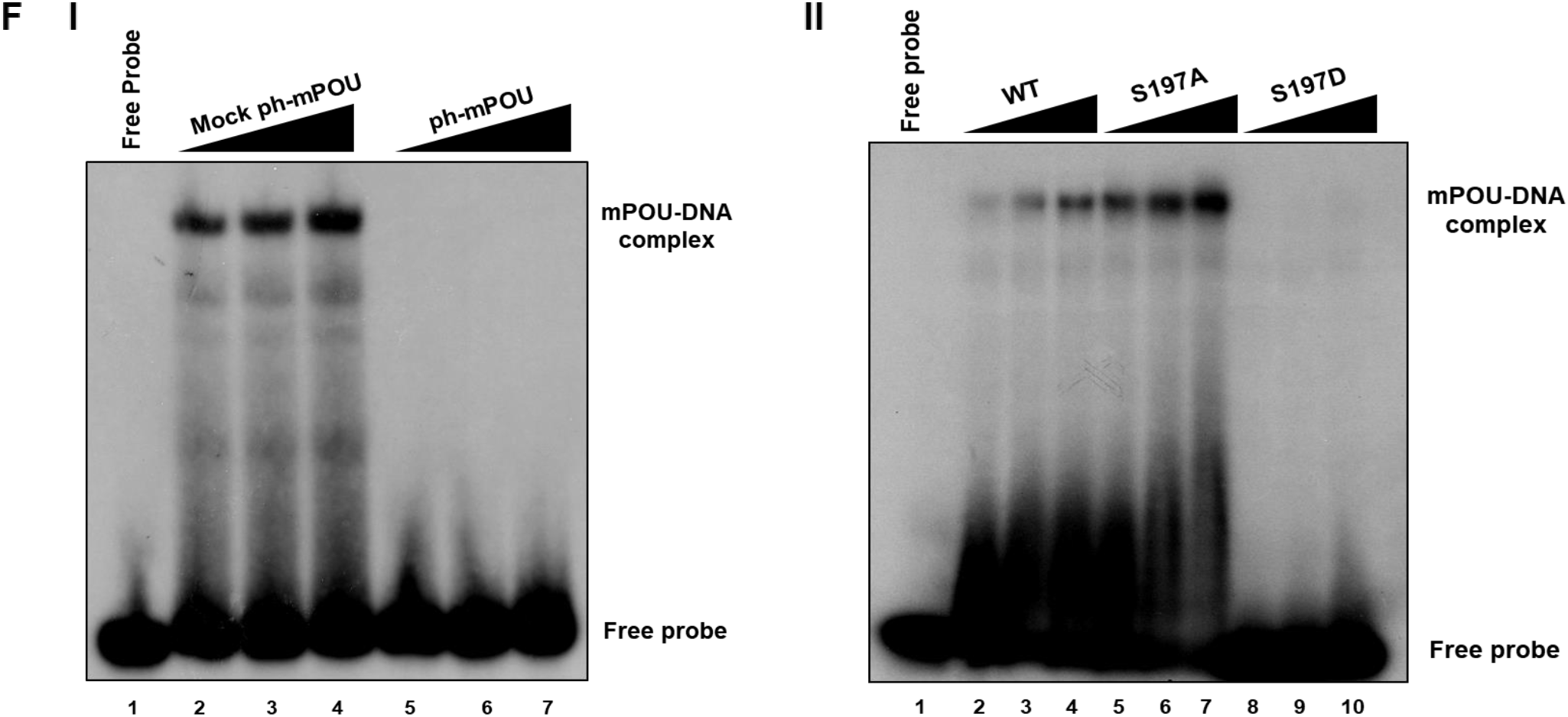
AurkA phosphorylates mPOU at S197 and inhibits its DNA binding. **A.** Schematic representation of mPOU domain organization. The N-terminal domain of the protein contains Gln and Pro-rich amino acids of unknown function. The DNA binding POU-specific (I) and POU-homeodomain (III) are joined together by a linker region (II). **B.** Western blot representation of differential migration pattern of Flag-mPOU protein from G1/S and G2/M synchronized cells. HEK293 cells were transiently transfected with Flag-mPOU plasmid constructs and synchronized at ether G1/S or G2/M phases by double thymidine block or nocodazole treatment respectively. Whole cell lysates prepared from these cells were separated by SDS-PAGE and probed with anti-Flag antibody for detection of Flag-mPOU. **C.** Identification of the major phosphorylation residue of mPOU by AurkA. His6-mPOU WT or the indicated point mutants, generated by site directed mutagenesis, were subjected to *in vitro* kinase assay using recombinant His6-AurkA purified from Sf21 insect cells using suitable baculoviral constructs. The reaction mix was separated by SDS-PAGE, stained by coomassie brilliant blue (CBB), dried and exposed to an X-ray film. The CBB staining for each of the lanes is shown below the autoradiogram profiles for comparing the loading levels across each lane. **D.** Colocalization of mPOU and γ-tubulin at the centrosome. HEK293 cells were stained for the indicated antibodies using indirect immunofluorescence. **E.** Colocalization of endogenous mPOU and AurkA. HEK293 cells were stained for the indicated antibodies using indirect immunofluorescence. Images were acquired at 63X magnification under oil immersion. Scale bar represent 5μm in length. **F.** Electrophoretic mobility shift assay using either recombinant His6-mPOU with (ph-mPOU) or without (mock ph-mPOU) prior *in vitro* phosphorylation by AurkA **(panel-I)** or using the WT and phosphorylation mutants (S197A/S197D) of His6-mPOU **(panel-II)** using a radiolabeled DNA oligonucleotide probe derived from the mPOU gene promoter.

To elucidate both the tissue-specific function of mPOU and the importance of its phosphorylation by Aurora kinase, we used C2C12 myoblasts as a differentiation model for *ex vivo* analyses. We find mPOU as a negative regulator of skeletal muscle differentiation. Furthermore, as our earlier study signify Aurora kinase A (AurkA) to be an enzyme necessary for maintenance of the differentiated state of skeletal muscle through E2F4 phosphorylation, our present results illuminate the multi-modal approach the kinase adopts through mPOU phosphorylation, to facilitate and maintain differentiation. Our findings suggest AurkA to be a quintessential player which governs the process of skeletal muscle differentiation.

## Materials and Methods

### Cell culture and reagents

HEK293 cells were cultured in DMEM (Sigma) with 10% fetal bovine serum (FBS). C2C12 myoblasts were maintained in DMEM containing 20% FBS (growth media; GM). Differentiation was induced by removing the GM, rinsing the cells with warm 1X phosphate buffered saline (PBS) and addition of DMEM containing 2% horse serum for various time periods. Different POU6F1 constructs were co-transfected into C2C12 myoblasts with pBabe-Puro vector and the cells were selected with puromycin, 24 hours post transfection, to enrich the transfected population, which were further used for differentiation experiments. For knockdown studies, C2C12 myoblasts were transfected with either control siRNA (Silencer™ Negative Control No. 1 siRNA; Invitrogen # AM4611) or a mixture of four siRNA pool targeting mPOU (ON-TARGET plus SMART pool; Dharmacon #19009), using RNAiMAX Reagent (Invitrogen™).

### Antibodies used

Various immunoblotting and immunofluorescence analyses were performed using anti-Flag (F1804; Millipore) and anti-β actin (A3854; Millipore), anti-myogenin (F5D; DSHB), anti-myosin heavy chain (MF20; DSHB), anti-POU6F1 (polyclonal antibody raised against a specific mPOU peptide; Abgenex), and anti-troponin T (T6277; Millipore).

### Cell synchronization

HEK293 cells, transiently transfected with Flag-mPOU, were synchronized in either G1/S or G2/M phases of the cell cycle by double thymidine block or thymidine-nocodazole treatment respectively. Briefly, 30-40% confluent asynchronously growing cells were treated with 2mM Thymidine for 18 hours (first block). The cells were then released from the block by washing twice with warm 1X PBS and then by addition of DMEM containing 10% FBS. 9 hours post release in complete media, 2 mM thymidine was again added to the cells and harvested 17 hours later. Mitotic (G2/M) block was similarly performed by addition of 2 mM thymidine for 24 hours, followed by washing twice with 1X PBS and releasing in DMEM containing 10%FBS for 3hrs. 100 ng/ml of nocodazole was finally added for 12 hours following which they were harvested for further analysis.

### Real time qPCR

Control or variously treated cells were harvested by trypsinization and lysed using TRIzol™ (Invitrogen™) and total RNA isolated by phenol: chloroform extraction followed by precipitation using isopropanol. The extracted RNA was reconstituted in diethyl pyrocarbonate (DEPC) treated water. cDNA strand was prepared using oligo-dT_23_ (Sigma), M-MLV Reverse Transcriptase (Sigma) and RNaseOUT™ Recombinant Ribonuclease Inhibitor (Invitrogen™) as per manufacturer’s recommendations. This cDNA was used for real time PCR (RT-PCR) analysis using SYBR-Premix Ex Taq™ II (Tli RNaseH Plus) (Takara) with 10 pmol of specific primers. RT-PCR reactions were carried out in Step One Plus™ Real-Time PCR (Applied Biosystems) machine and amplification protocols were followed as indicated in the manufacturer’s protocol. Fold expression change was calculated using ΔΔCt method using actin gene primers as internal control. The primer sequences for the various qPCR primers used are depicted in Table 1.

**Table 1.**
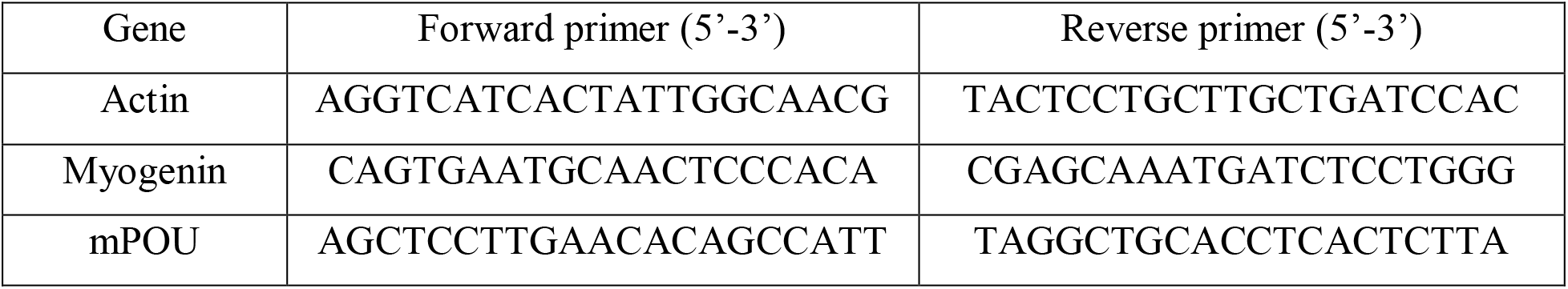

### *In vitro* kinase assay

Various recombinant proteins were expressed in *E. coli* and incubated with His6-tagged Aurora kinase A, expressed in Sf21 cells using suitable baculoviral constructs, in a 30 μl reaction mixture containing 50 mM tris-HCl, 100 mM NaCl, 0.1 mM EGTA, 10 mM MgCl_2_, 0.2% β-mercaptoethanol and [γP^32^] ATP (Specific Activity 3.5 Ci/mmol). The reaction mixture was incubated at 30°C for 15 minutes. The reaction was inhibited over ice for 5mins, constituent proteins were denatured by the addition of loading dye containing SDS, heated at 90°C for 5mins and resolved using 12% denaturing PAGE, followed by autoradiography using X-ray films. For mass phosphorylation, 1μg of mPOU WT was incubated with AurkA at 30°C for 30 mins in the presence of 1mM ATP. This was followed by three replenishments with the same amount of enzyme and ATP at every half hour interval. The final incubation was prolonged for 2 hours. As control, mock phosphorylation reactions were carried out with the same components except the enzyme under similar conditions. The reaction mix were separated on a 12% SDS PAGE, followed by autoradiography using X-ray films.

### Electrophoretic mobility shift assay

40pmol of single stranded oligonucleotide was labelled at the free 5’-OH group with 20 units of T4 polynucleotide kinase (NEB # M0201) in a 50 μl reaction volume containing 1X polynucleotide kinase buffer and 30 μCi γ^32^P ATP at 37°C for 30min. At the end of the reaction 150 μl of water was added and subjected to phenol: chloroform extraction. The extracted aqueous layer containing the labelled oligonucleotide was precipitated along with 10 μg of tRNA by using ethanol and sodium acetate and then dried and dissolved in annealing buffer containing 10 mM tris-HCl, and 20 mM NaCl at pH 8. To create the double stranded probe, the radiolabeled single strand of the Brn5 promoter (5’-GATCTGCTCCTGCATGCCTAATAGG-3’) was allowed to anneal with the complementary unlabeled oligonucleotide (3’-CTAGACGAGGACGTACGGATTATCC-5’) in annealing buffer containing 10 mM tris-HCl and 20 mM NaCl at pH-8, by first heating the oligo-mix to 100°C and then allowing the temperature to gradually drop to room. The double stranded oligo is then purified by native-PAGE and extracted in 1X tris-EDTA (TE) buffer (10 mM Tris-HCl, pH 8 and 1 mM EDTA) at 37°C. This was further purified by gel filtration using C40 packed syringe column. The probe was finally eluted in TE buffer and diluted to 5000 cpm/μl and allowed to form complex with the DNA binding protein of interest in appropriate binding buffer for 30 minutes at recommended temperature (25-30°C). The resultant complex was resolved on a native-PAGE and then subjected to autoradiography to visualize the complex and its mobility shift.

### Bromodeoxyuridine (BrdU) incorporation assay

The proliferative capacity of cells was measured using a BrdU cell proliferation assay (Roche, USA) according to the manufacturer’s instructions. Briefly, C2C12 myoblasts expressing various mPOU constructs were incubated with BrdU labelling solution for 30 min and fixed in 4% paraformaldehyde. These were subsequently stained with monoclonal antibody against BrdU for one hour, followed by FITC conjugated secondary antibody and imaged using a fluorescent microscope at 20X magnification.

### Chimera generated 3D-model

The crystal structure of mPOU (or Brn5)-DNA complex was downloaded from Protein Data Bank (PDB ID: 3D1N) and Chimera software was empoyed to generate the three-dimensional image, highlighting the various residues.

## Results

### mPOU is phosphorylated by Aurora kinases and inhibits its DNA binding ability

To investigate whether mPOU harbors differential post translational modifications in a cell cycle dependent manner, we studied the migration pattern of transiently transfected Flag-mPOU in cells synchronized in different phases of the cell cycle. Upon blocking cells in the G2/M phase, Flag-mPOU displayed retarded migration as compared to the G1/S blocked cells (Fig.1B).

Several residues of mPOU were found to be phosphorylated by AurkA in the *in vitro* high-throughput screening (7). We chose some of the high scoring Ser/Thr sites, namely S183, S197, T239 and S240, and mutated them to Ala by site directed mutagenesis. The respective *E. coli* purified phospho-deficient mutants were then subjected to *in vitro* kinase assay to identify the major phosphorylation site(s). Intriguingly, S197A phospho-deficient mPOU mutant exhibited the maximal reduction in phosphorylation by AurkA, as compared to the other point mutants (Fig.1C). In order to confirm the existence of S197 as a phosphorylated residue in cells, we transfected Flag-mPOU transiently into either HeLa cells or C2C12 myoblasts and found the existence of phosphorylation of S197 in both the cell lines (Supplementary Fig. 1).

We next sought to investigate the localization of endogenous mPOU in mitotic HEK293 cells and observed mPOU to be resident at the centrosomes (Fig.1D). Furthermore, the mPOU also co-localized with AurkA (Fig.1E). These results collectively indicate a probable phosphorylation of mPOU by AurkA at the onset of mitosis, specifically at the centrosomes.

Ser197 is a crucial residue within the POU-specific domain of mPOU and forms hydrogen bonding with the phosphate groups of the DNA backbone (10). It was, therefore, imperative to hypothesize if phosphorylation of this amino acid can disrupt the DNA contact, due to an ionic repulsion and hence, the subsequent DNA binding. Concordantly, we observed that *in vitro* phosphorylation of the recombinant His6-mPOU protein by AurkA led to a complete loss of the canonical DNA binding ability of WT-mPOU, as observed by gel shift assay (Fig.1F; panel-I). Similar result was also observed for the phospho-mimic (S197D) mPOU mutant (Fig. 1F; panel-II), thus strengthening the possibility of charge-repulsion in the abrogation of DNA contact by phosphorylated mPOU.

### Negative role of mPOU on muscle differentiation is mitigated by AurkA mediated phosphorylation

mPOU displays tissue-restricted expression patterns during mice development. While the expression is exclusively brain specific during embryogenesis, it is more cluttered in brain, heart, skeletal muscle and lungs in the adults (13). In order to gain insights into the function of mPOU, we focused on adult skeletal myogenesis as a physiologically relevant scenario in the context of mPOU expression and probable function and chose C2C12 myoblasts as a model system. We observed that mPOU was downregulated upon induction of differentiation (Fig.2A). Sustained overexpression of Flag-mPOU WT inhibited (Fig.2B), while knockdown of endogenous mPOU enhanced differentiation of the C2C12 myoblasts, as assessed by differentiation markers (Fig.2C; panel-II). These results signify mPOU as a negative regulator of myogenic differentiation and also highlight the necessity of mPOU downregulation in order to catalyze the differentiate process.

**Figure 2:**
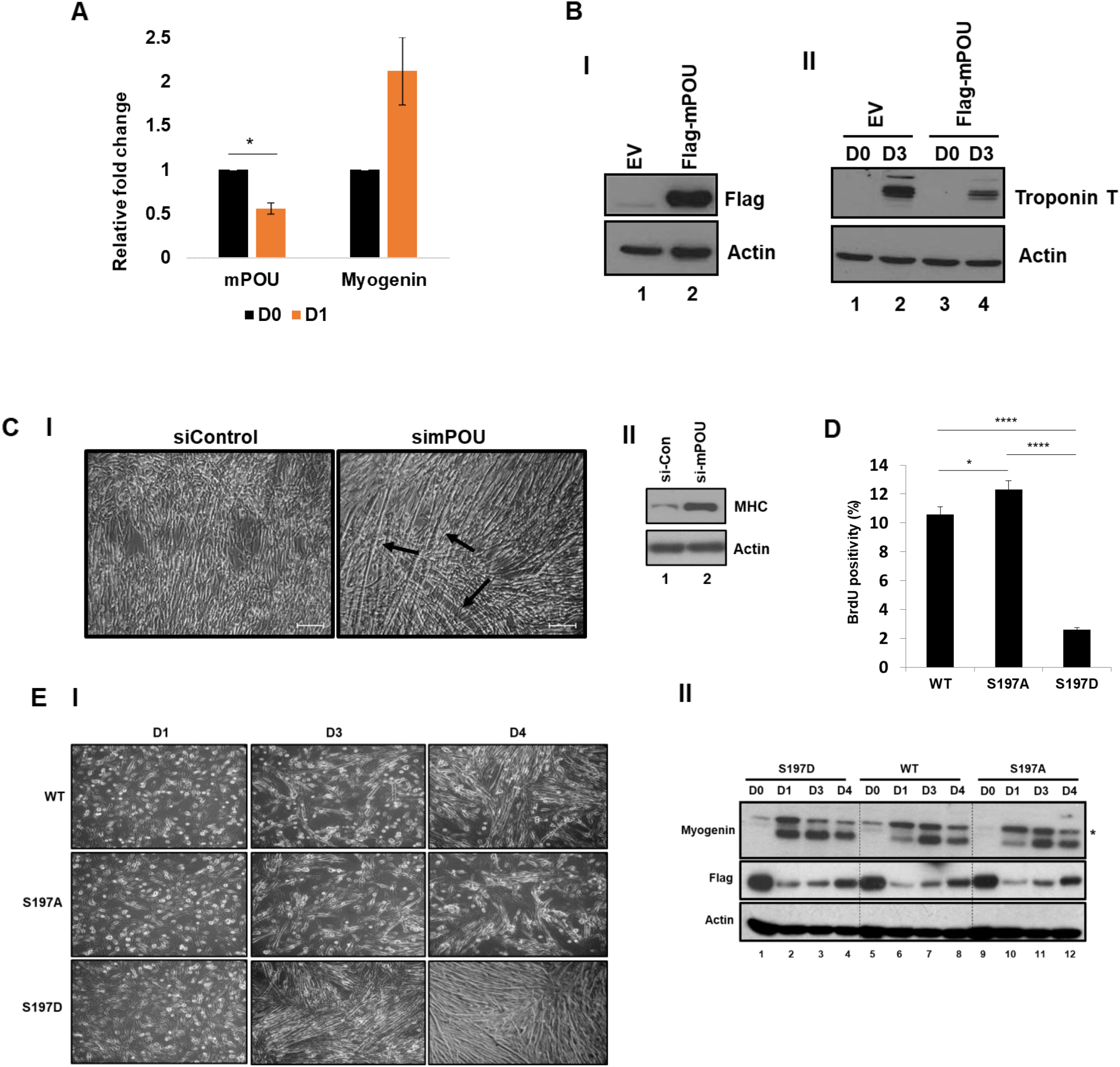
AurkA mediated mPOU phosphorylation is necessary for myogenic differentiation. **A.** mPOU is downregulated upon induction of C2C12 differentiation. Real-time qPCR results indicating the expression of the mentioned genes. Myogenin was considered as a positive control. Unpaired Student’s t-test was carried out to carry out statistical analysis of the samples. Error bars represent standard deviation from two independent biological replicates. *p<0.05. **B.** Western blot representation of Flag-mPOU overexpression in C2C12 myoblasts **(panel-I)** and subsequent effect on day3 post-differentiation induction, studied by the indicated antibodies **(panel-II). C.** Bright field images representing the morphology of C2C12 myoblasts upon control and mPOU siRNA treatments, 48 hours post induction of differentiation. Images were acquired at 10X magnification. Scale bar represent 20μm in length. **(panel-I).** Western blot representation for expression of differentiation specific protein marker (myosin heavy chain; MHC) expression under control siRNA and mPOU siRNA treatment conditions **(panel-II). D.** mPOU WT and phosphorylation mutants exhibit altered BrdU incorporation capabilities. C2C12 myoblasts were transfected with the indicated Flag fusion constructs of mPOU and subsequently incubated with BrdU and stained with anti-BrdU monoclonal antibody. The degree of proliferation was quantified as % BrdU positive cells and plotted as a bar chart (n=300). Unpaired Student’s t-test was carried out to carry out statistical analysis of the samples. Error bars represent standard deviation from two independent biological replicates. *p<0.05, ****p<0.0001. **E.** Bright field images representing the morphology of C2C12 myoblasts upon overexpression of the indicated Flag-mPOU constructs and upon induction of differentiation for the indicated time periods. Images were acquired at 10X magnification **(panel-I).** Western blot representation for expression of differentiation specific protein marker (myogenin) for the indicated Flag-mPOU constructs and the indicated time periods **(panel-II).**

Myogenic differentiation necessitates exit from active cellular cycling and therefore, inhibits further proliferation. As our results indicate that mPOU acts as a negative regulator of the differentiation state, we studied the potency of either WT or phospho-mutants of mPOU in arresting proliferation, through BrdU incorporation assay. Strikingly, reduction of C2C12 proliferation for the phospho-mimic (S197D) mutant, while not for the phospho-deficient (S197A) mutant or the WT, was observed (Fig.2D). We next asked whether the diminished cellular proliferation for S197D mutant can prime the cells for differentiation and observed that the S197D mutant displayed enhanced differentiation (Fig.2E). These results comprehensively suggest that mPOU phosphorylation by AurkA primes myoblasts for differentiation through an inhibition of proliferation.

## Discussion

The present study broadens the functional repertoire of phosphorylation dependent regulation of skeletal muscle differentiation by Aurora kinase A (AurkA), through a tissue-restricted transcription factor, POU6F1 (or mPOU). We found Ser197 (S197) to be the major site of phosphorylation of mPOU by AurkA. Intriguingly, S197 directly contacts the sugar-phosphate backbone of the cognate DNA (10) and therefore, may pose a major hindrance to DNA binding. This presumption was corroborated by mobility shift assay and we observed that either *in vitro* AurkA phosphorylated mPOU WT or phospho-mimic S197D (SD) mutant mPOU protein lacked the ability to bind its own promoter DNA sequence. Indirect immunocytochemistry of cells revealed an important corelate for the localization of mPOU at the centrosomes, and further corroborated by colocalization with AurkA at the mitotic phase. We conjecture that dwelling of mPOU at the centrosome may drive its phosphorylation by AurkA, which in turn primes its exclusion from the early mitotic DNA. It is possible that dephosphorylation of mPOU upon mitotic exit, enable it to re-associate with the chromatin for transcription. Interestingly, mPOU possess a distinct RVXF sequence at its C-terminus which represents a well-conserved PP1 binding motif (15). Three-dimensional overview suggests a close proximity between the RVXF sequence and the S197 residue (Supplementary Fig.2) and therefore, strengthens the chance of mPOU dephosphorylation by PP1 in the later mitotic phases or upon mitotic exit.

In order to investigate the tissue specific roles of mPOU and the physiological relevance of AurkA dependent mPOU phosphorylation, we exploited C2C12 myoblasts as a surrogate for skeletal muscle tissue where mPOU expression is reported earlier (13). We found that mPOU knockdown enhanced while its overexpression retarded the differentiation process. We also found that WT or phospho-deficient mutant inhibited while the phospho-mimic mutant exhibited enhanced differentiation. We conclude mPOU as a negative regulator of skeletal muscle differentiation and envisage AurkA mediated mPOU phosphorylation as a trigger for the loss of mPOU occupancy onto its own gene promoter which in turn, results into its transcriptional downregulation and facilitates a switch to a heightened differentiation induction (Fig.3).

**Figure 3:**
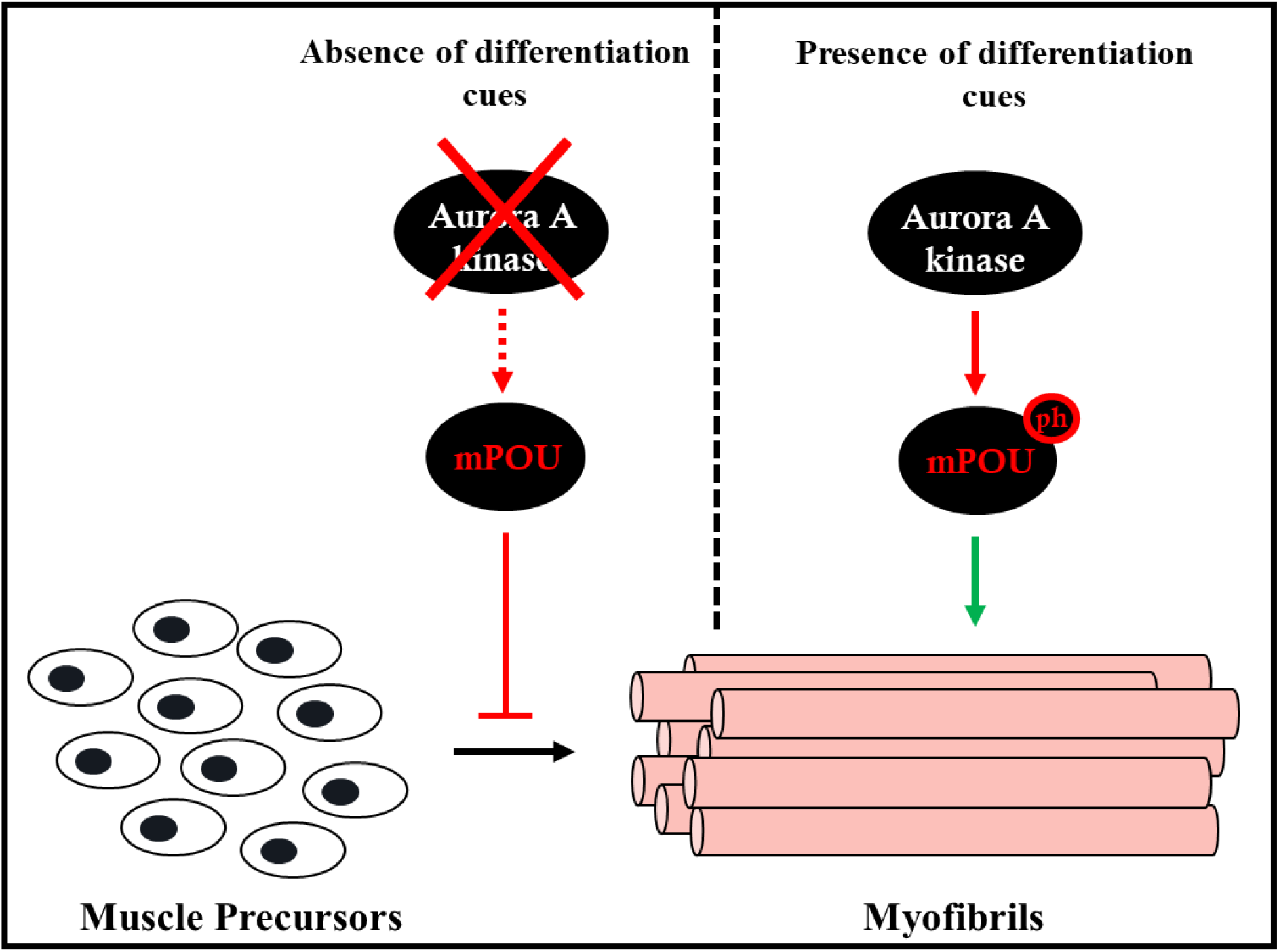
Model depicting the role of AurkA in mediating skeletal muscle differentiation though mPOU phosphorylation and DNA binding inhibition. Differentiation signals trigger AurkA activation which phosphorylates mPOU and inhibits its DNA binding ability. This event causes reduced mPOU gene transcription, which otherwise act as a negative modulator of differentiation, and facilitates the process.

AurkA serve as an important regulator of centrosome maturation and in determining the extent of microtubule assembly at the onset of mitosis. Various signals emanating from the Golgi leads to the enhanced recruitment and phosphorylation of AurkA at the centrosomes (16,17). The role of calcium dependent signaling is also pivotal in AurkA activation dynamics (18,19). Although calcium dependent events drive muscle differentiation (20–23), whether calcium signaling has a role to play in muscle differentiation through AurkA activation presents an interesting avenue for future studies. It is feasible that an injury driven stimulus activates calcium signaling, which in turn potentiates AurkA activity and subsequent cascade of mPOU phosphorylation to drive the process of differentiation (Fig.3).

## Acknowledgements

The authors thank Suma Bangalore Srinivas and Mr. Prajwal, Jawaharlal Nehru Centre for Advanced Scientific Research (JNCASR) for confocal imaging. K.D. and A.B. were supported by grants from Government of India (GOI)’s Council of Scientific and Industrial Research (CSIR). T.K.K. acknowledges JNCASR for partial financial support to V.J.R. T.K.K. is a recipient of Sir J.C. Bose National Fellowship, Department of Science and Technology, GOI. This work was also supported by grants from the Department of Biotechnology, GOI, through Chromatin and Disease: Program Support (Grant BT/01/ CEIB/10/111/01 dated 30.09.2011) and by Indo-Japan bilateral joint research program by Department of Science and Technology, India, and Japan Society for the Promotion of Science (JSPS), Japan. Studies at Tohoku University was supported in part by grants-in-aid from JSPS (17K07278 and 25670156).

---

**Figure S1:**
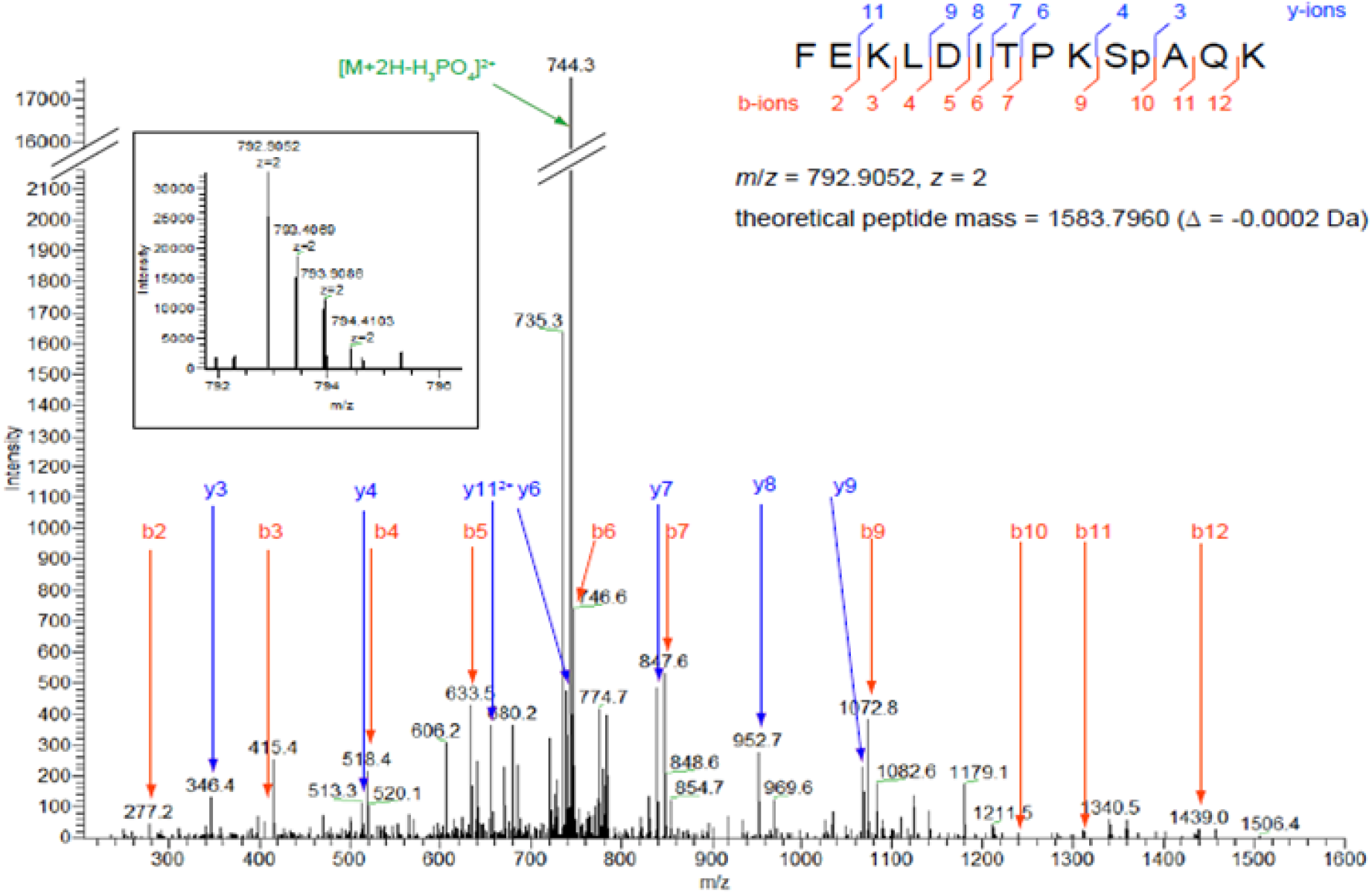
Representative mass spectra of S197 phoshorylation of mPOU in HeLa cells.

**Figure S2:**
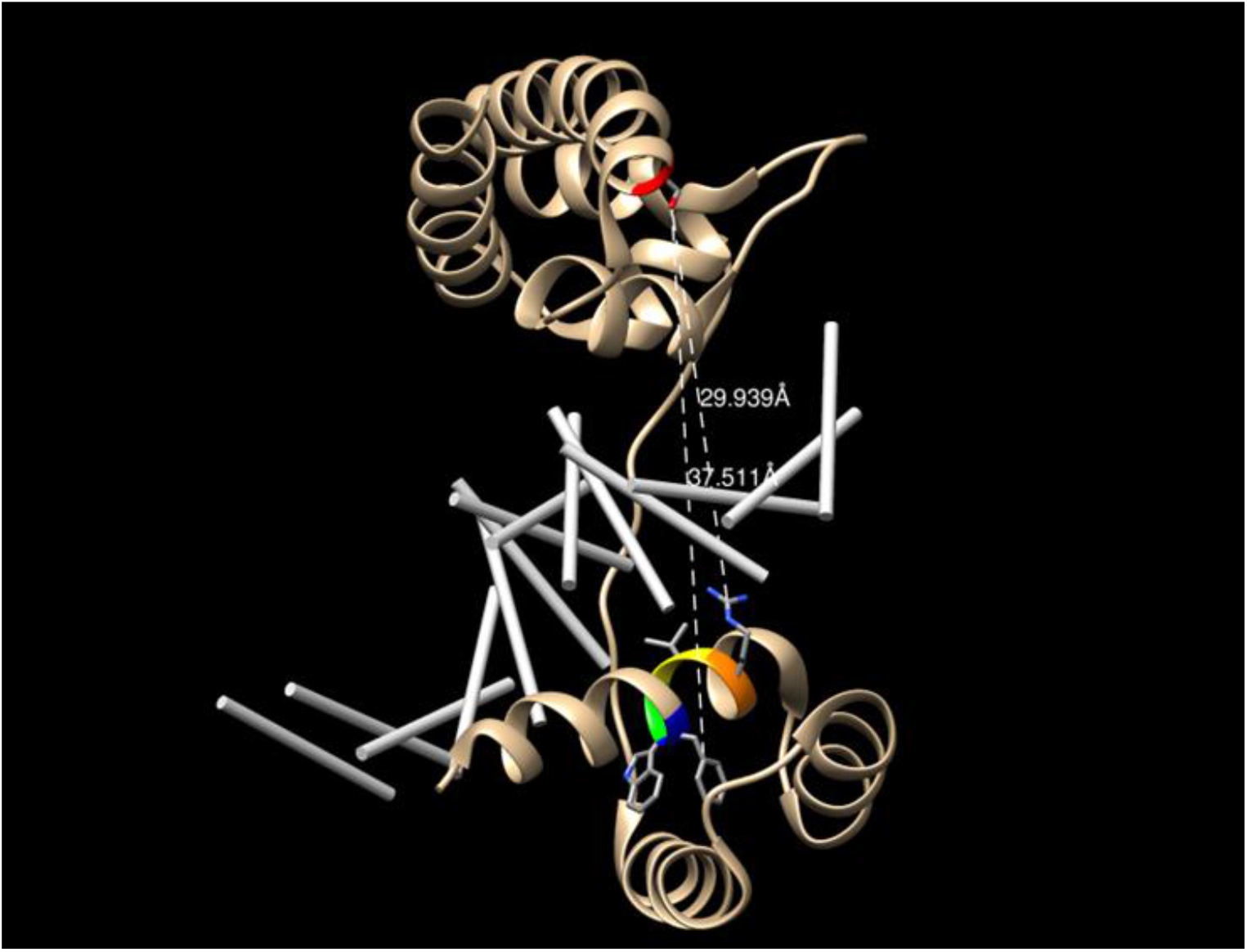
Chimera generated three-dimensional spatial overview of the proximity of S197 site to the RVWF (PP1 recognition) motif. Distance between serine 197 (S197, red) and the PP1 recognition RVWF motif (Arginine, orange; Valine, yellow; Tryptophan, green; Phenylalanine, blue) measured in Å (PDB ID: 3D1N).

## References

1. Dhawan J, Rando TA. (2005) Stem cells in postnatal myogenesis: molecular mechanisms of satellite cell quiescence, activation and replenishment. Trends Cell Biol. 15(12):666–73.

2. Weintraub H (1993) The MyoD family and myogenesis: Redundancy, networks, and thresholds. Cell 75:1241–1244.

3. Walsh K, Perlman H (1997) Cell cycle exit upon myogenic differentiation. Curr Opin Genet Dev. 7:597–602.

4. Planas-Silva MD, Weinberg RA (1997) The restriction point and control of cell proliferation. Curr Opin Cell Biol 9:768–772.

5. Mal A, et al. (2000) p21 and retinoblastoma protein control the absence of DNA replication in terminally differentiated muscle cells. J Cell Biol. 149:281–292.

6. Amabile G1, D’Alise AM, Iovino M, Jones P, Santaguida S, Musacchio A, Taylor S, Cortese R. (2009) The Aurora B kinase activity is required for the maintenance of the differentiated state of murine myoblasts. Cell Death Differ. 16(2):321–30.

7. Dhanasekaran K, Bose A, Rao VJ, Boopathi R, Shankar SR, Rao VK, Swaminathan A, Vasudevan M, Taneja R, Kundu TK. (2019) Unraveling the role of aurora A beyond centrosomes and spindle assembly: implications in muscle differentiation. FASEB J. 33(1):219–230.

8. R.A. Sturm, W. Herr. (1988) The POU domain is a bipartite DNA-binding structure. Nature, 336, pp. 601–604

9. W. Herr, M.A. Cleary. (1995) The POU domain: versatility in transcriptional regulation by a flexible two-in-one DNA-binding domain. Genes Dev., 9, pp. 1679–1693

10. Pereira JH, Kim SH. (2009) Structure of human Brn-5 transcription factor in complex with CRH gene promoter. J Struct Biol. 167(2):159–65.

11. Lakich MM, Diagana TT, North DL, Whalen RG. (1998) MEF-2 and Oct-1 bind to two homologous promoter sequence elements and participate in the expression of a skeletal muscle-specific gene. J Biol Chem. 273(24):15217–26

12. Ryan AK, Rosenfeld MG. (1997) POU domain family values: flexibility, partnerships, and developmental codes. Genes Dev. 11(10):1207–25

13. Wey E, Lyons GE, Schäfer BW. (1994) A human POU domain gene, mPOU, is expressed in developing brain and specific adult tissues. Eur J Biochem. 220(3):753–62.

14. Wey E, Schäfer BW. (1996) Identification of novel DNA binding sites recognized by the transcription factor mPOU (POU6F1). Biochem Biophys Res Commun. 220(2):274–9.

15. Wakula P, Beullens M, Ceulemans H, Stalmans W, Bollen M. (2003) Degeneracy and function of the ubiquitous RVXF motif that mediates binding to protein phosphatase-1. J Biol Chem. 278(21):18817–23.

16. Persico A, Cervigni RI, Barretta ML, Corda D, Colanzi A. Golgi partitioning controls mitotic entry through Aurora-A kinase. (2010) Mol Biol Cell. 21(21):3708–21.

17. Barretta ML, Spano D, D’Ambrosio C, Cervigni RI, Scaloni A, Corda D, Colanzi A. (2016) Aurora-A recruitment and centrosomal maturation are regulated by a Golgi-activated pool of Src during G2. Nat Commun. 7:11727.

18. Plotnikova OV, Pugacheva EN, Dunbrack RL, Golemis EA. (2010) Rapid calcium-dependent activation of Aurora-A kinase. Nat Commun. 1:64.

19. Plotnikova OV, Pugacheva EN, Golemis EA. (2011) Aurora A kinase activity influences calcium signaling in kidney cells. J Cell Biol. 193(6):1021–32. doi:10.1083/jcb.201012061.

20. Porter GA Jr, Makuck RF, Rivkees SA. (2002) Reduction in intracellular calcium levels inhibits myoblast differentiation. J Biol Chem. 277(32):28942–7.

21. Benavides Damm T, Egli M. (2014) Calcium’s role in mechano transduction during muscle development. Cell Physiol Biochem. 33(2):249–72.

22. Friday BB, Horsley V, Pavlath GK. (2000) Calcineurin activity is required for the initiation of skeletal muscle differentiation. J Cell Biol. 149(3):657–66.

23. Nasipak BT, Padilla-Benavides T, Green KM, Leszyk JD, Mao W, Konda S, Sif S, Shaffer SA, Ohkawa Y, Imbalzano AN. (2015) Opposing calcium-dependent signalling pathways control skeletal muscle differentiation by regulating a chromatin remodelling enzyme Nat Commun. 6:7441.

